# Usp16 modulates Wnt signaling in primary tissues through Cdkn2a regulation

**DOI:** 10.1101/326272

**Authors:** Maddalena Adorno, Benedetta Nicolis di Robilant, Shaheen Sikandar, Veronica Haro Acosta, Jane Antony, Craig Heller, Michael F. Clarke

**Affiliations:** Institute for Stem Cell Biology and Regenerative Medicine, Stanford University School of Medicine, Stanford, California 94305, USA; Molecular and Computational Biology Department, University of Southern California, Los Angeles, California 90087 USA; Department of Biology, Stanford University School of Medicine, Stanford, California 94305, USA

## Abstract

Regulation of the Wnt pathway in stem cells and primary tissues is still poorly understood. Here we report that Usp16, a negative regulator of Bmi1/PRC1 function, modulates the Wnt pathway in mammary epithelia, primary human fibroblasts and MEFs, affecting their expansion and self-renewal potential. In mammary glands, reduced levels of Usp16 increase tissue responsiveness to Wnt, resulting in upregulation of the downstream Wnt target Axin2, expansion of the basal compartment and increased *in vitro* and *in vivo* epithelial regeneration. Usp16 regulation of the Wnt pathway in mouse and human tissues is at least in part mediated by activation of Cdkn2a, a regulator of senescence. At the molecular level, Usp16 affects Rspo-mediated phosphorylation of LRP6. In Down’s Syndrome (DS), triplication of Usp16 dampens the activation of the Wnt pathway. Usp16 copy number normalization restores normal Wnt activation in Ts65Dn mice models. Genetic upregulation of the Wnt pathway in Ts65Dn mice rescues the proliferation defect observed in mammary epithelial cells. All together, these findings link important stem cell regulators like Bmi1/Usp16 and Cdkn2a to Wnt signaling, and have implications for designing therapies for conditions, like DS, aging or degenerative diseases, where the Wnt pathway is hampered.

## INTRODUCTION

Wnt signaling has a crucial role in the normal function of several stem cell types, including mammary, neural and embryonic stem cells^1, 2^. Wnt is also very tightly regulated during aging, and, in the majority of tissues, Wnt signaling declines during senescence^3, 4^ Furthermore, the decline of Wnt signaling with age contributes to the pathogenesis of osteoporosis^5^, Alzheimer’s disease, and Parkinson’s disease^6^. However, despite several decades of studies focusing on this pathway, its regulation in primary tissues, especially stem cells, remains only partially understood.

Interestingly, the Wnt decline during aging parallels an increase in levels of p16^Ink4a,^ a protein coded at the *Cdkn2a* locus^7–9^. The *Cdkn2a* locus is tightly regulated by USP16 and by Bmi1, a member of the Polycomb Repressive Complex 1 (PRC1). USP16 is a deubiquitination enzyme that plays a crucial role in regulating tissue homeostasis and stem cell self-renewal and expansion^10^. USP16 acts by detaching a monoubiquitin protein from histone H2A-K119, opposing the epigenetic repressive function of PRC1^11^. Bmi1 is a member of the PRC1 complex and a crucial regulator of stem cell self-renewal in several adult tissues, including the bone marrow and the brain^12, 13^. Together, Bmi1/PRC1 and USP16 provide a robust and elaborate mechanism regulating the epigenetic landscape of stem cells, and governing the equilibrium between self-renewal and senescence^10^.

Here we show an unexpected link between Wnt signaling and Bmi1/USP16, connecting two important signaling pathways acting on stem cells and primary tissues. We find that USP16 acts as a negative regulator of Wnt signaling, and that its action is mediated at least in part by the Bmi1/USP16 regulated target *Cdkn2a.*

## RESULTS

### MMTV-Wnt1-Usp16^+/-^ preneoplastic mammary epithelial cells are more proliferative then their wild-type counterparts

We first investigated mammary tissue, where the Wnt pathway plays a pivotal role in orchestrating proper mammary gland development and stem cell maintenance^14^ As we previously reported, increased levels of Usp16 inhibit normal mammary gland stem cell function

10. We therefore decided to investigate the effect of Usp16 dosage levels in MMTV-Wnt1 mouse mammary glands. MMTV-Wnt1 transgenic mice present ectopic activation of Wnt1, causing a powerful mitogenic effect on the mammary epithelium. As a consequence, these animals develop extensive ductal hyperplasia early in life^15^. The ratio between basal and luminal cells more than doubles in MMTV-Wnt1 animals compared to wild-type animals (Fig. 1A and Suppl. Fig.S1A), as observed by Fluorescence Activated Cell Sorting (FACS) analyses. This increase in basal cells is in line with previous studies^16^, for example evidences from LRP5 mutant mice in which a decrease in the activation of the Wnt/β-catenin reduces the percentage of cells within the basal compartment of the mammary gland^17^ Preneoplastic mammary glands derived from MMTV-Wnt1 mice crossed with Usp16^+/-^ mice showed that the basal expansion was significantly higher in MMTV-Wnt1-Usp16^+/^” glands (almost 2-fold over the basal/luminal ratio observed in MMTV-Wnt animals) (P=0.0073) (Figure 1A), suggesting Usp16 might have a role in limiting the activation of the Wnt pathway. Histological analyses also showed an increase in the number (P<0.0001) and the area (P=0.0071) of ducts present in the breast tissue of 3-months old virgin females (Figure 1B-C). To further investigate the effects of Wnt activation in Usp16^+/-^ mammary cells, we performed *in vitro* colony formation plating breast epithelial cells sorted based on the expression of EpCAM, CD49f, and lineage markers (CD31, CD45 and Ter119) (Suppl. Figure S1B). Cells were plated on a feeder layer of murine cells producing Wnt3a ligand that sustains *in vitro* long term expansion of mammary progenitors^18^. MMTV-Wnt1-Usp16^+/-^ cells generate more than twice as many colonies compared to MMTV-Wnt cells after the first passage (Figure 1D) (P<0.001). Taken together, these data show that Usp16 limits the activation of the Wnt pathway in mammary epithelials, affecting the growth of basal cells.

**Figure 1.**
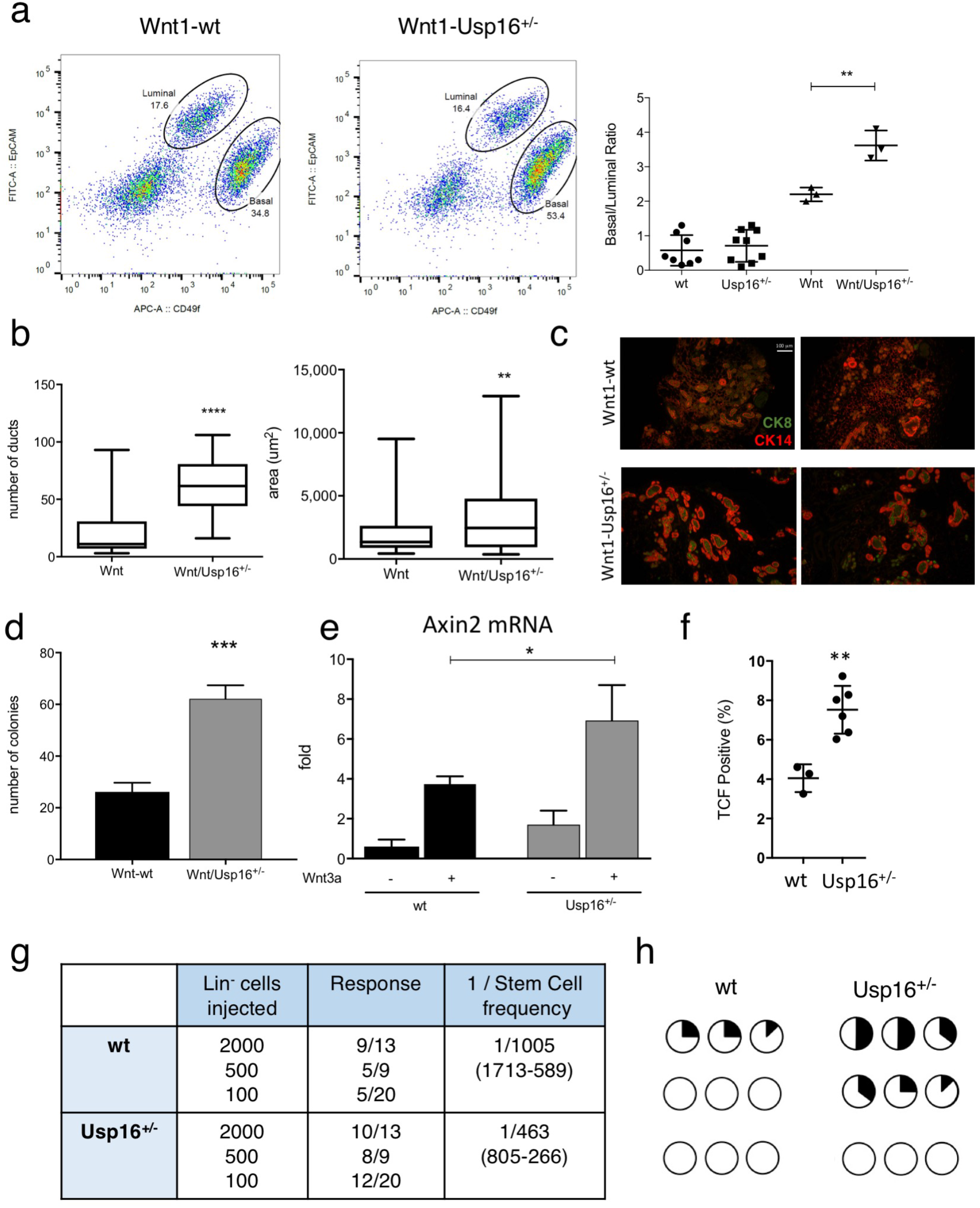
Heterozygosis of Usp16 in mammary tissue promotes Wnt-driven *in vitro* and *in vivo* cell expansion. (**A**) FACS analysis shows a higher basal to luminal cell ratio in the preneoplastic mammary gland of virgin MMTV-Wnt1-Usp16^+/-^ mice. On the left, representative FACS plots of Lin^−^ (Ter119^−^ CD45^−^ CD31^−^) mammary cells for the indicated genotypes. On the right, quantification of basal/luminal cell ratio. Each dot represents an individual mouse. (**B-C**) Histological analyses of preneoplastic mammary glands reveal an increase in the number and area of ducts derived from MMTV-Wnt1-Usp16^+/-^ mice. The graph shows the average of six slides analyzed per animal, two animals per group. Quantification was performed with ImageJ software. On panel C, two representative pictures per genotype are shown. Keratin 8 and Keratin 14 were used to mark luminal and basal cell layers, respectively. The white bar scale is 100 μm. (**D**) FACS-sorted epithelial cells from MMTV-Wnt1-Usp16^+/-^ preneoplastic mammary glands form more colonies *in vitro* compared to control mice (n=3 per group). Shown is passage P1. (**E**) Usp16^+/-^ sorted mammary epithelial cells show an increased induction of Axin2 mRNA levels 16 hours after Wnt3A stimulation (50ng/ml). Three independent experiments were performed. (**F**) The mammary epithelial TCF-GFP^+^ frequency is increased in Usp16^+/-^ compared to wt TCF-GFP animals after one passage *in vitro.* Each dot represents an individual animal. Unpaired T-test shows a two-tailed pvalue <0.01. (**G**) Limiting dilution analysis (ELDA) shows an increased frequency of mammary repopulating cells in Usp16^+/-^ compared to wt breast tissue. Four rounds of transplants were performed. The table shows the stem cell frequency and the 95% confidence interval between lower and upper values. (**H**) Usp16^+/-^ cells were able to engraft better than wt cells in secondary recipients. Two independent experiments were performed. Each circle represents a recipient. In black, the estimated area occupied by the mammary tree in the fat pad two months after transplantation.

### Usp16^+/-^ mammary cells are more responsive to Wnt3a activation and show a higher engraftment potential

To evaluate the interaction of Usp16 and the Wnt pathway in a more physiological context, we compared the growth of epithelial mammary cells derived from wild-type and Usp16^+/-^ mice when cultured directly on Matrigel in the presence of 50 ng/ml of Wnt3a ligand. After one passage, Usp16^+/-^ cells generated almost 50% more colonies than cells with two copies of Usp16 (P<0.05) (Suppl. Figure S2A). To investigate the molecular activation of the Wnt pathway, we FACS-sorted mammary epithelial cells from wild-type and Usp16^+/-^ mice and looked at the expression of Axin2, a canonical transcriptional target of the Wnt pathway containing eight Tcf/LEF consensus binding sites in its promoter^19^. Activation of Axin2 mRNA was almost doubled in Usp16^+/-^ cells compared to their wild-type counterpart (P=0.0115) (Figure 1E). T_o_ further understand if the role of Usp16 was limited only to a subset of mammary cells, we sorted basal and luminal cells based on the expression of CD49f and EpCAM^20^. Basal cells were very responsive to Wnt3a as measured by induction of Axin2, while luminal cells showed lower levels of activation (Suppl. Figure S2B). However, both basal and luminal cells heterozygous for Usp16 showed higher expression of Axin2 in response to Wnt3a if compared to wild-type cells (in basal cells approximatively 40-fold versus 17-fold, respectively) (P<0.01) (Suppl. Figure S2B). To further assess the regulation of the Wnt pathway by Usp16, we took advantage of TCF/LEF-H2B/GFP mice, a reporter model that allows tracking of Wnt-expressing cells through GFP^21^. In line with previous reports, the presence of GFP cells in the mammary gland was very limited^22^ (Suppl. Figure S2C). However, after one passage of *in vitro* culture, the observed frequency of GFP+ epithelial cells increase from 4% in wild-type cells to 8% in Usp16^+/-^ cells (P<0.01) (Figure 1F and Suppl. Figure S2D).

Since mammary gland epithelial cells from Usp16^+/-^ mice have increased Wnt activation and given the role of Wnt in mammary epithelial cell expansion, we hypothesized that mammary gland epithelial cells from the Usp16^+/-^ mice would have increased *in vivo* reconstitution ability. To test this hypothesis, serial dilution transplantation assays of wild-type and Usp16^+/-^ mammary cells in cleared fat pads of syngeneic FVB mice were performed. Analyses of 42 recipient mice revealed a greater then 2-fold increase in the frequency of Usp16^+/-^ mammary cells able to reconstitute a fat pad compared to wild-type cells (stem cell frequency: 1/463 in Usp16^+/-^ cells compared to 1/1005 in wt cells) (P<0.05) (Figure 1G and Suppl. Figure S2E). We also performed secondary transplantation, an essay that well correlates with the true replication capacity of cells 16. After transplantation of 500 cells, six out of nine recipients showed a mammary tree generated by Usp16^+/-^ donor cells. Conversely, engraftment in only three out of nine recipients of wt cells was observed (Figure 1H).

Taken together, these data show that dosage of Usp16 regulates the activation of the Wnt pathway in the breast tissue, and that Usp16 haploinsufficiency increases the *in vitro* and *in vivo* expansion potential of mammary progenitor cells.

### *Cdkn2a* acts as a negative regulator of Wnt activation in murine and human cells

Usp16 controls H2A ubiquitination levels at the Cdkn2a locus^10^. We therefore wondered if *Cdkn2a* could itself be a regulator of the Wnt pathway. Similar to what we observed in MMTV-Wnt1-USP16^+/-^ mice, FACS analyses of the mammary tissue revealed an increase in the basal/luminal ratio in three-months old *Cdkn2a*^+/-^ mice compared to wild-type mice (P<0.05) (Figure 2A and Suppl. Figure S3A). Moreover, real-time PCR analyses revealed an increase in expression levels of Axin2 in *Cdkn2a^+/-^* basal cells, compared to cells derived from age-matched wild-type mice (P<0.05) (Figure 2B).

**Figure 2.**
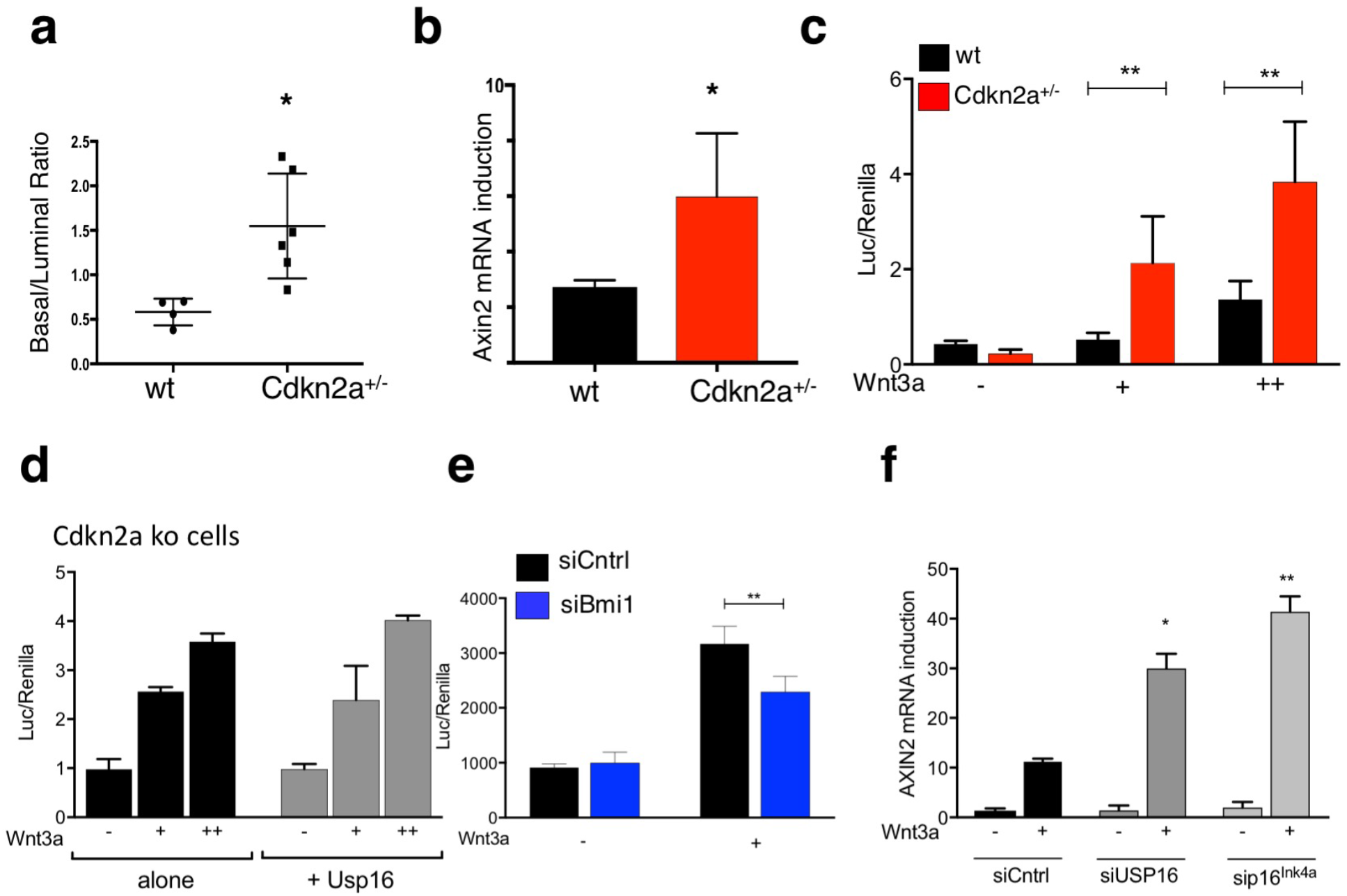
Cdkn2a mediates Usp16 response to Wnt3a. (**A**) Basal cell expansion is observed in the mammary gland of Cdkn2a^+/-^ mice. A Graph quantifying the basal/luminal ratio is shown. Each dot represents an individual mouse. (**B**) Cdkn2a^+/-^ sorted basal cells show an increased induction of Axin2 mRNA levels after 16 hours of Wnt3A stimulation (50ng/ml). Two independent experiments were performed for a total of 4 animals per group. (**C**) Terminal Tip Fibroblasts (TTFs) derived from Cdkn2a^+/-^ animals show an increase response to two different doses of Wnt3a (20 ng/ml and 50 ng/ml). The graph shows activation of a Top/Flash reporter for Wnt activity normalized by Renilla expression, cotransfected along with the reporter. (**D**) TTFs derived from Cdkn2a^−/-^ animals and transfected with a vector expressing a Usp16 transgene or a control vector were analyzed for their ability to activate the Top-flash reporter. No differences were observed with two different doses of Wnt3a (20 ng/ml and 100 ng/ml). (**E**) MEFs cells were transiently transfected with two individual siRNAs targeting Bmi1. Activation of 6KD Wnt reporter is shown, normalized by Renilla expression. Cells were treated with 100 ng/ml of Wnt3a for 16 hours. (**F**) Two different human foreskin cultures were transiently transfected with siRNA targeting USP16, p16^Ink4a^ or a scramble sequence. USP16 and p16^Ink4a^ downregulation promotes the induction of AXIN2 mRNA expression 24 hours after treatment with 100 ng/ml of Wnt3a.

To further evaluate the role of *Cdkn2a* in the regulation of the Wnt pathway, Terminal Tip Fibroblasts (TTFs) were derived from *Cdkn2a^+/-^, Cdkn2a^−/-^*, and matching control animals. Cotransfection of a luciferase βcatenin/TCF reporter 6xKD^23^ with a plasmid expressing Renilla was performed in TTFs at passage 2. Stimulation with increasing doses of Wnt3a activated the luciferase reporter, and the levels of activation were significantly higher in *Cdkn2a^+/-^* cells at every tested dose (P<0.01) (Figure 2C). A similar trend was observed, as seen in breast tissue, in TTFs derived from Usp16^+/-^ mice (P<0.0001) (Suppl. Figure S3B). To test the epistatic relationship between Usp16 and *Cdkn2a,* transfection of a plasmid expressing Usp16 was performed in *Cdkn2a^−/-^* TTFs. The 6xKD reporter was activated similarly in response to increasing doses of Wnt3a both in the presence and in the absence of Usp16 overexpression, suggesting that Cdkn2a is a downstream effector of Usp16 on the regulation of the Wnt pathway (Figure 2D). Similar results were observed in HEK293T human cells (Suppl. Figure S3C); HEK293T cells, as the majority of cell lines, do not have an intact *Cdkn2a* signaling pathway^24, 25^. The *cdkn2a* locus codes for two proteins: p16^Ink4a^ and p14^Arf^. In particular p16^Ink4a^ expression levels have been correlated with cellular senescence and aging^7, 8^. Mouse embryonic fibroblasts (MEFs) at early passage showed increased activation of the Wnt reporter 6xKD in the presence of siRNAs targeting Usp16 or p16^Ink4a^ (P<0.01) (Suppl. Figure S4A and S4B), in line with our previous results in TTFs. Noticeably, the same cells, if treated with two different siRNAs targeting Bmi1, which inhibits Cdkn2a expression, as predicted showed decreased Wnt response (Figure 2E and Suppl. Figure S4A) (P<0.01).

Finally, to assess the biological relevance of the connection between Usp16 and Cdkn2a in human tissues, we used two different lines of human primary fibroblasts at early passages. After transient delivery of interfering-RNAs targeting USP16 and CDKN2a (Suppl. Figure S4B), we noticed that downregulation of USP16 increased AXIN2 mRNA induction three-fold compared to cells transfected with a control siRNA (P<0.05) (Figure 2F). The same effect was triggered by p16^Ink4a^ downregulation (P<0.01)(Figure 2F).

Taken together, these data show that the *Cdkn2a* locus is crucial for the Usp16/PRC1-mediated control of the Wnt pathway in murine and human primary cells.

### Usp16 and p16^Ink4a^ modulate Rspo-mediated LRP6 phosphorylation

The Wnt pathway can be modulated via multiple mechanisms including expression of different extracellular receptor components as well as variations in the expression of intracellular signal transducers. To further elucidate how Usp16 impacts Wnt activation, a direct link between Usp16 and βcatenin was tested. Immunoprecipitation studies in MEF cells did not show any physical interaction between the two proteins (Suppl. Figure S5A). However, silencing of USP16 or the cdkn2a coded p16^Ink4a^ in MEF cells (Suppl. Figure S4A) increased levels of Wnt-induced βcatenin nuclear accumulation (Fig 3A). Therefore, Usp16 and p16^Ink4a^ are able to affect the Wnt pathway upstream of βcatenin.

**Figure 3.**
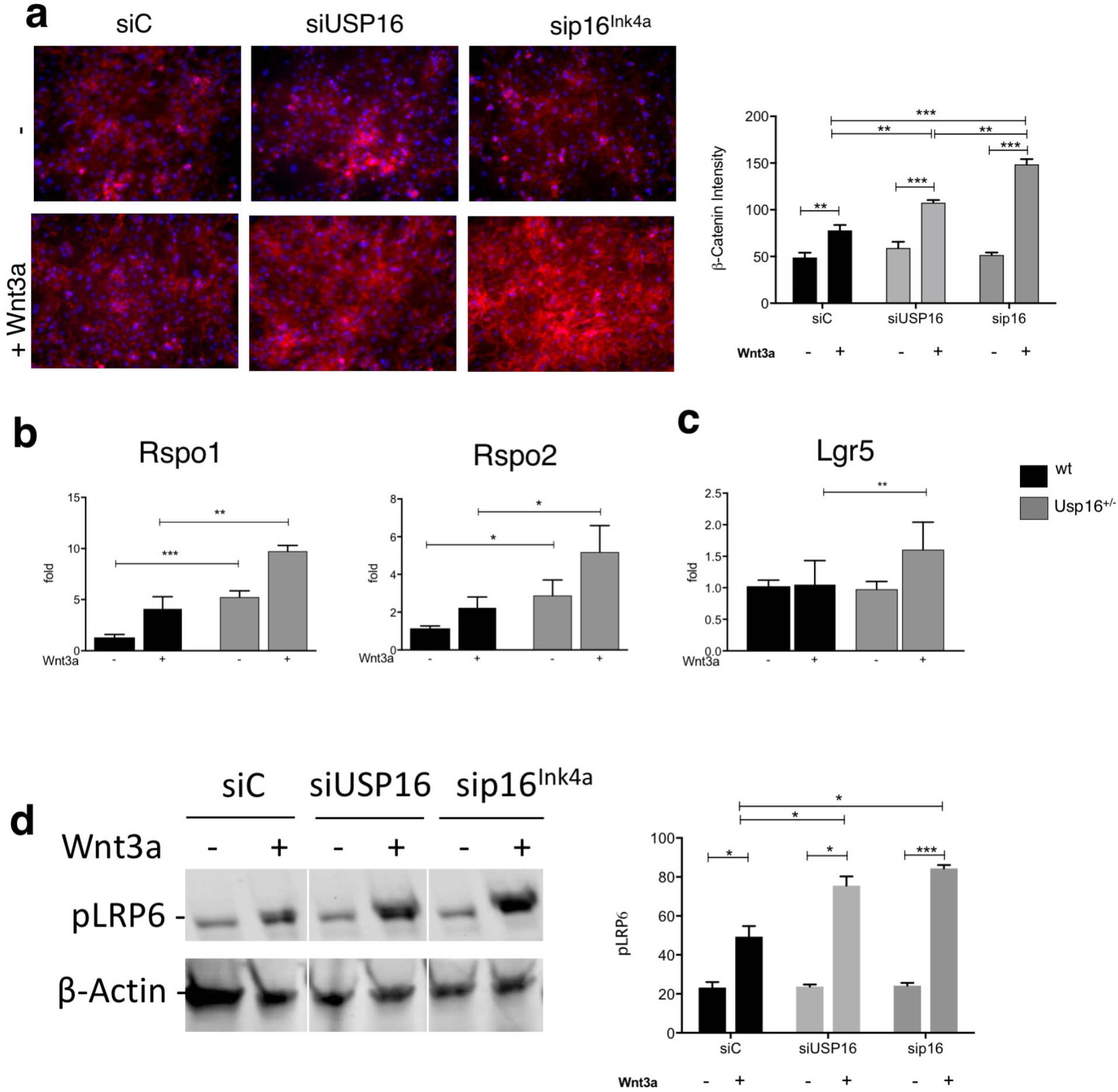
Usp16 and p16^Ink4a^ modulates Rspo-mediated LRP6 phosphorylation. (A) MEFs cells were transiently transfected with siRNAs targeting Usp16 or p16^Ink4a^ and treated with or without Wnt. Representative staining for βcatenin (in red) and DAPI (in blue) is shown on the left. On the right, the graph shows quantification of βcatenin intensity. Treatment with siRNA against Usp16 or p16^Ink4a^ increases levels of βcatenin upon treatment with 50 ng/ml of Wnt3a. Experiments were repeated twice with two different MEF cultures. (**B**) Usp16^+/-^ mammary epithelial cells, both untreated and treated with Wnt3a, express higher levels of Rspo1 and Rspo2 (n=3 per group). (**C**) FACS-sorted mammary basal cells heterozygous for Usp16 express higher levels of Lgr5 upon treatment with Wnt3a (n=4 per group). (**D**) Extracts derived from MEF cells and treated with the indicated siRNA were tested by western blot. Phospho-LRP6 immunoblots are shown on the left panel. βactin is used as loading control. A white line has been introduced between samples for clarity, but the image has not been cropped. On the right, the graph shows quantification of phospho-LRP6 intensity. Experiments were repeated four times.

Rspo1 and Rspo2 are potent growth factors for many adult stem cells including those of the mammary gland^26, 27^ They enhance Wnt signaling through interaction with their receptor, Lgr4/5/6, potentiating LRP phosphorylation^28, 29^. Rspo1/2 were expressed at higher level in breast cells heterozygous for Usp16 both with or without Wnt3a treatment (P<0.01 for Rspo1, P<0.05 for Rspo2) (Figure 3B). Moreover, Wnt3a-treated breast basal cell express higher levels of Lgr5 compared to their wild-type counterparts (P<0.01) (Figure 3C).

Next, levels of LRP6 phosphorylation were tested. Western blot studies showed that silencing of USP16 or p16^Ink4a^ increases Wnt-induced levels of LRP6 phosphorylation (Figure 3D). This is not further increased by Rspondin treatment (250ng/ml). LRP6 phosphorylation is dependent on the presence of Rspondin, as proven by concomitant treatment of MEFs with siRNA targeting Usp16 (or cdkn2a) and Rspo1/2 (Suppl. Fig. S4A and S5B).

Taken together, these data show that Usp16 and p16^Ink4a^ act mostly upstream of the βcatenin complex, through the regulation of the Rspondin/Lgr5/LRP6 axis.

### Trisomy of Usp16 affects the Wnt pathway in Ts65Dn cells and human DS-derived cells

Activation of the Wnt pathway is altered in a number of conditions: upregulated in many types of cancers, and downregulated in bone diseases, metabolic diseases and aging^1^. As a proof of principle, we wanted to investigate if the accelerated aging and decreased self-renewal observed in Down’s syndrome (DS) and previously associated with trisomy of Usp16^10^ was also associated with aberrant Wnt signaling. To test this, we first took advantage of BAT-GAL transgenic mice, a reporter strain that expresses β-galactosidase in the presence of activated βcatenin, mimicking the pattern of Wnt signaling^30^. We crossed these mice with Ts65Dn mice, the most widely used model for Down’s Syndrome^31, 32^. Ts65Dn mice are trisomic for 132 genes homologous to genes on human chromosome 21, including Usp16.

We FACS-sorted mammary epithelial cells from wt and Ts65Dn reporter mice based on the expression of CD24, CD49f, and lineage markers^16^. These cells were tested by real time PCR for expression of β-gal RNA. Ts65Dn-derived cells showed a dramatic decrease in expression of the Wnt-activated reporter (P<0.01) (Fig 4A). To assess whether Usp16 plays a role in Wnt modulation in the mouse DS model, we derived TTFs from wt, Ts65Dn and Ts65Dn/Usp16^+/-^ mice. The Ts65Dn/Usp16^+/-^ mice are only normalized for Usp16 copy number, and not for the other genes triplicated in this model. Co-transfection of a luciferase β-catenin/TCF reporter (TOP-FLASH)^33^ with a plasmid expressing Renilla showed a decreased Wnt activation in Ts65Dn-derived fibroblasts (P<0.0001), while normalization of Usp16 copy number reverted this effect to normal levels (P=0.0021)(Figure 4B).

**Figure 4.**
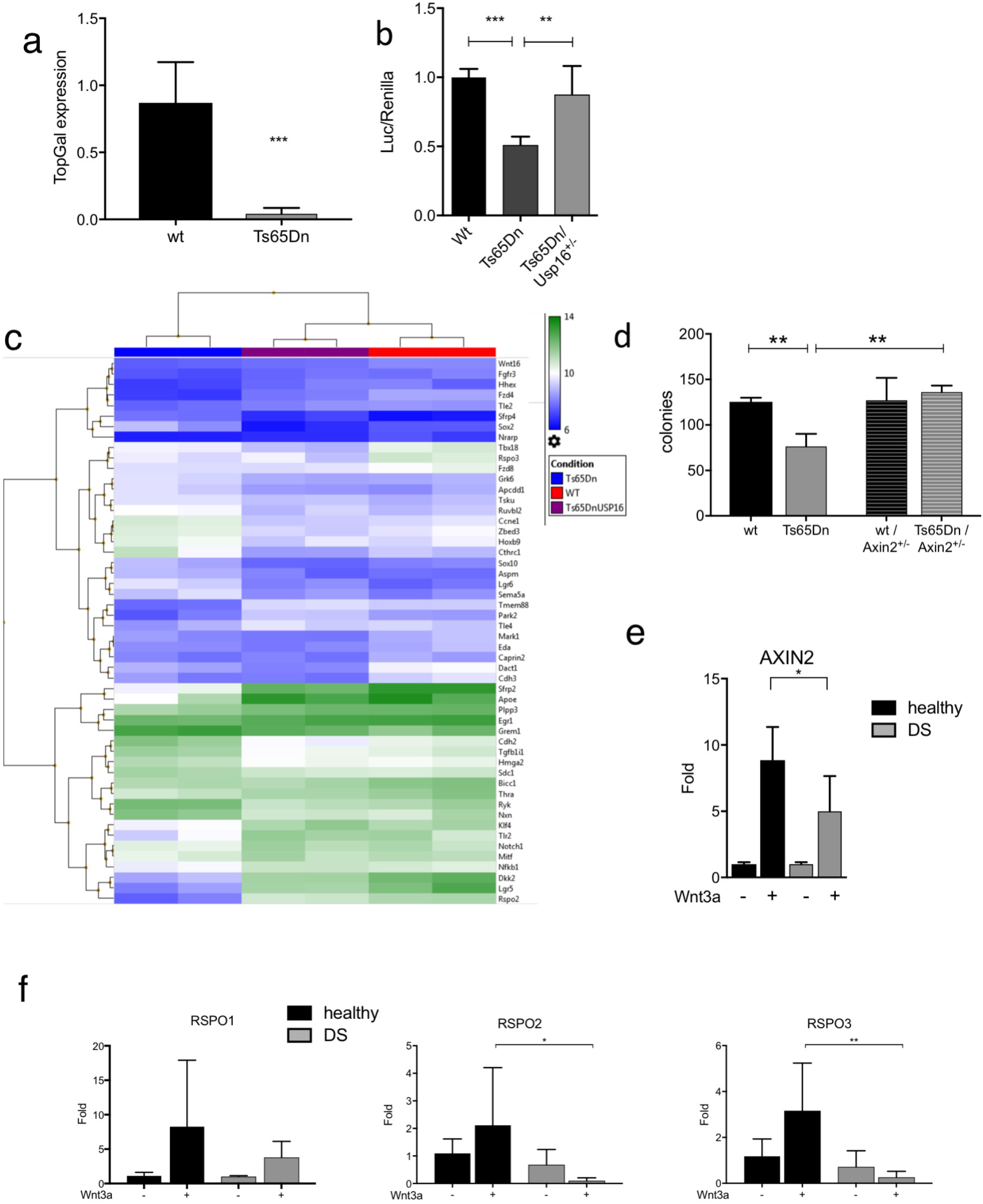
Usp16 trisomy affects the Wnt pathway in Down’s Syndrome. (**A**) SYBR Green analyses revealed a marked decrease in the expression of a Top-Gal reporter of Wnt activity in mammary epithelial cells FACS-sorted from Ts65Dn mice compared to wt animals (n=3 per group). (**B**) TTFs derived from animals with the indicated genotypes were analyzed for their ability to activate the Top-flash reporter in response to 20 ng/ml of Wnt3a. Data were normalized based on induction in wt animals. Wt and Ts65Dn/Usp16^+/-^ cells activate Top-Flash more efficiently than Ts65Dn-derived cells. (wt, n=5; Ts65Dn, n=5; Ts65Dn/Usp16^+/-^, n=2). (**C**) Microarray analysis shows clustering of wt, Ts65Dn and Ts65Dn/Usp16^+/-^ RNA expression based on a Wnt signature of genes differentially expressed between wt and Ts65Dn cells (P<0.05). (n=2 per group). Ts65Dn/Usp16^+/-^ RNA expression (samples in the middle) clusters with wt expression (samples on the right). (**D**) FACS-sorted epithelial cells from Ts65Dn mammary glands form less colonies *in vitro* compared to their wt counterpart. Axin2 heterozygosis increases the ability of Ts65Dn cells to form colonies (n=3 animals per group; n=2 for wt animals). Shown is passage P0. (**E,F**) Real time PCR analyses showed that human DS foreskin fibroblasts are less responsive to Wnt3a than cells derived from healthy patients in the activation of AXIN2, RSPO2 and RSPO3 (healthy, n=4: DS, n=3).

To further elucidate Usp16-derived pathway regulation, we performed microarray analyses on these TTFs. Using TAC software, we performed cluster-analyses of these samples (F-test <0.001 for wt vs Ts65Dn). Notably, we found that Ts65Dn/Usp16^+/-^ cells cluster with wild-type and not with Ts65Dn cells, suggesting that one single gene, namely Usp16, is largely responsible for the aberrant activation of the Wnt pathway that we reported in Down’s syndrome cells (Suppl. Fig. S6). We then analyzed the expression pattern of a series of genes associated with the Wnt pathway (the list was derived from the gene ontology annotations of MGI, see Suppl. Table T1).

The clustering shows again that normalization of Usp16 copy number reverts Ts65Dn expression to a pattern very similar to the one observed in wild-type cells (F-test <0.05 for wt vs Ts65Dn, see Suppl. Table T2) (Figure 4C). Notably, Rspo2, Rspo3 and Lgr5 are regulated by Usp16 in Ts65Dn cells.

We also wanted to see if upregulation of the Wnt pathway was able to rescue the *in vitro* colony forming ability of Ts65Dn mammary epithelial cells. We crossed Ts65Dn mice with Axin2^−/-^ mice, obtaining a progeny expressing only one copy of Axin2, a negative regulator of the Wnt pathway^34^ As previously shown^10^, Ts65Dn epithelial mammary cells generated approximately half as many colonies as wild type cells when plated on a feeder layer providing Wnt3a. However, heterozygosis of Axin2 rescues this defect (P<0.01) (Figure 4D).

To assess whether the Wnt pathway was affected in human DS tissues as well, we tested foreskin fibroblasts derived from Down’s syndrome and control individuals. We analyzed by Real-time PCR the activation of the canonical TCF/LEF target AXIN2 in response to treatment with 50 ug/ml of Wnt3a. Wnt3a drove a robust upregulation of this gene in tissues derived from healthy donors, but the expression was not as strong in trisomic cells (P<0.05) (Figure 4E). Given the role played by Usp16 in regulating R-spondins in the mammary tissue, we tested their expression in these cells as well. Wnt3a-driven upregulation of Rspo2 and Rspo3 was completely lost in DS cells (P<0.01 for Rspo2, P<0.001 for Rspo3) (Figure 4F), suggesting a conserved mechanism for Usp16-mediated regulation of Wnt pathway across species. Rspo1 mRNA was also detected but we didn’t notice a significant difference in pattern of expression.

Taken together, these data support the idea that trisomy of Usp16 plays a role in modulating the Wnt pathway in the context of Down’s syndrome, and that Wnt modulation can improve the defects observed in trisomic mammary gland and fibroblast cells.

## DISCUSSION

Our data show that Bmi1 and Usp16, important chromatin regulators in stem cells, modulate Wnt signaling in primary mammary epithelial and fibroblast cells. Levels of Usp16 affect the *in vitro* and *in vivo* expansion capabilities of mammary cells. This process is mediated by *Cdkn2a,* a well known biomarker for senescence^7–9^. To our knowledge, this is the first report highlighting a connection between the Bmi1/Usp16/Cdkn2a and Wnt pathways (Figure 5A). Usp16^+/-^ is associated with an increased expression of R-spondin and Lgr5, as well as an increase in levels of LRP6 phosphorylation, contributing to the amplification of Wnt signaling.

**Figure 5.**
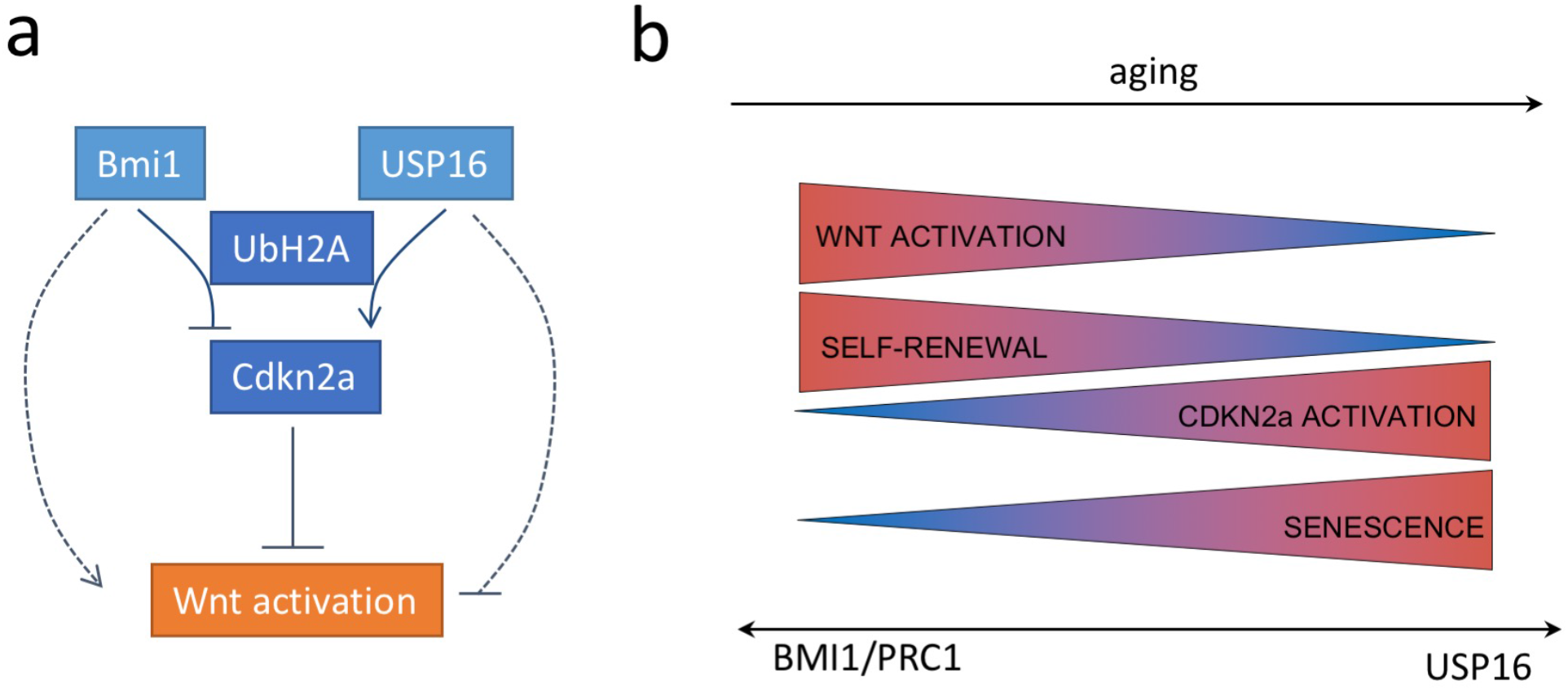
General mechanism of Wnt induction. (**A**) Proposed mechanism of interaction between Usp16 and the Wnt pathway. Bmi-1 and Usp16 control the level of ubiquitination of histone H2A. The balance between these two antagonistic proteins determines the levels of activation of *Cdkn2a. Cdkn2a* acts as a negative regulator of the Wnt pathway. Other mechanisms linking Bmi1 and Usp16 to Wnt activation are also possible. (**B**) During aging, increased tissue senescence and increased expression levels of p16^INK4a^ have been observed. At the same time, there is a concurrent decline in stem cell self renewal and Wnt activation. These events are correlated to a change in balance between Bmi1 - promoting selfrenewal and Wnt activation - and Usp16 - promoting cellular senescence and *Cdkn2a* expression.

It is widely reported that Wnt plays a crucial role in stem cell maintenance, aging and cellular senescence in several tissues, including the breast and the brain^3^. In general, canonical Wnt signaling is decreased in senescent cells and senescence can be delayed by providing extracellular canonical Wnt ligands^4^ However, we have limited knowledge of Wnt regulation in primary tissue and stem cells. The majority of studies aiming at describing the molecular regulation of Wnt signaling are based on ectopic expression of Wnt components in immortalized cell lines such as HEK293T cells^35–38^. Because these cells have undergone mutations such as inactivation of the Cdkn2a/Rb pathways, most of the signals related to cellular senescence are invariably disrupted in these cells. Thus, physiological Wnt regulators that require intact cellular pathways may not be observed in such cells. In this study, we were able to identify Usp16 and Cdk2na as important and previously unknown physiological regulators of Wnt signaling in primary fibroblasts and mammary epithelial cells (Figure 5B).

In this article we also report that trisomy of Usp16 in Down’s Syndrome is linked to a reduction in Wnt pathway activation. Interestingly, microarray analyses in TTFs show that Ts65Dn RNA expression profile, and in particular Wnt-related genes, are strongly affected by trisomy of Usp16. Down’s syndrome is associated with several signs of early aging, including early onset and higher incidence of Alzheimer’s disease and higher than expected rate of osteoporosis and diabetes^39, 40^. Our current work suggests that, at least in part, Usp16 may exert its role on senescence by modulating the Wnt pathway.

The results of our study have important implications for understanding the molecular regulation of development, aging, stem cell biology and cancer. They also offer proof of principle for a possible therapeutic approach to several conditions where Wnt signaling is altered. Modulation of Usp16 by small molecules could offer a potentially successful intervention for targeting several of these pathological conditions via downregulation of *Cdkn2a* and/or upregulation of the Wnt pathway.

## MATHERIALS AND METHODS

### Mice

B6SJL-Tg(Wnt1)1Hev/J (Stock number: 002870), Tg(TCF/Lef1 -HIST1H2BB/EGFP) 61Hadj/J (Stock number: 013752), B6.Cg-Tg(BAT-lacZ)3Picc/J (Stock number: 005317), Ts65Dn (stock 004850), B6N.129P2-Axin2tm1Wbm/J (Stock number: 009120) and FVB/NJ (Stock number: 001800) were purchased from Jackson Laboratories. Usp16^+/-^ mice (FVB/N-Usp16Tg(Tyr)2414FOve/Mmjax) were ordered from MMRRC. Mice were genotyped by realtime or by PCR as previously published^10^(Jackson website). p16^Ink4a-/-^p19^Arf-/-^ mice (B6.129-Cdkn2a^tm1Rdp^) were obtained from Mouse Models of Human Cancers Consortium (NCI-Frederick). Control littermates were used as wild type mice. Mice were housed in accordance with the guidelines of Institutional Animal Care Use Committee at Stanford Animal Facility. All experimental protocols were approved according to APLAC #10868.

### Mouse breast analysis and in vitro colony forming assays

Mouse mammary glands were dissected and analyzed as previously described^16^. Briefly, the glands were digested with collagenase/hyaluronidase for 2 hours followed by ACK. Then a 2 minutes Trypsin and DNAse/Dispase digestion was performed. The cells were then stained with the following antibodies: EpCAM-FITC, CD49f-APC, CD24-PE-Cy7 and Linage-PacBlue (CD45, Ter119 and CD31) (Biolegend). Viable cells were identified based on forward- and side-scatter profiles and by DAPI exclusion. Single cells were analyzed and sorted using FACS Aria II (BD Bioscience). For in vitro colony forming assay, 10.000 sorted epithelial cells were plated into 96-well plate previously coated with 2.5% growth factor reduced matrigel. If not otherwise specified, irradiated L1-Wnt3a expressing cells were used as feeder layer. Cells were grown into liquid media supplemented with Y-27632 dihydrochloride (Sigma-Aldrich), murine recombinant EGF (Prepotech), rhR-Spondin1 (R&D) and Nogging (R&D), as previously described^10^.

### Primary and secondary mammary transplants

Lineage^−^ (CD45^−^CD31^−^Ter119^−^) cell populations were isolated from 12-week mice in staining media and resuspended in 10μl of sterile PBS + 30% matrigel per transplant before being injected into the cleared fat pads of 21-24 day old recipient FVB/NJ mice as previously described^16^. Frequency of long-term reconstituting cells from limiting dilution experiments was calculated using ELDA software^41^. For secondary mammary transplants, mammary glands were digested and 500 linage^−^ cells injected in secondary FVB/NJ recipients. All transplants were allowed to grow for at least 8 weeks but not more than 10 weeks before analysis. For mammary transplant outgrowth area calculation, NIH Image J software was used. Briefly, mammary ducts were measured with the free-hand tool by drawing a shape around the duct. Measurements were performed in a ‘blind’ fashion and at the same magnification for all samples. The entire fat pad was used to determine the maximum area coverage (100%). Only positive outgrowths were used in the measurement.

### Immunofluorescence of mammary tissue

MMTV-Wnt1 and MMTV-Wnt1Usp16^+/-^ 12-week old mice were euthanized and mammary glands were surgically removed. Glands were fixed in formalin overnight and then transferred to 70% ethanol. They were then embedded in paraffin and sectioned for histology. For staining the slides were deparafinised in xylene and alcohol grades. Antigen retrieval was carried out in Tris-EDTA buffer by heating in a microwave for 20min. Primary antibodies CK14 (Covance) and CK8 were applied overnight. Secondary antibodies were anti-rat DyLight 488 and anti-rabbit DyLight 594 (both from Jackson Labs). Sections were then mounted using Prolong Anti-fade reagent (Invitrogen). Images were taken with a NIKON inverted microscope. Area and duct calculations were done using NIH Image J software.

### RNA expression analysis

For real-time analyses, RNA was extracted using RNeasy Plus Micro or Mini Kit (Qiagen) and complementary DNA obtained using SuperScript Vilo cDNA Synthesis (Invitrogen). When less than 10,000 cells were sorted, pre-amplification using TaqMan PreAmp (Applied Biosystems) was performed. Real-time reactions were assembled using Taqman probes (Applied Biosystems) in accordance with the manufacturer’s directions. Expression data were normalized by the expression of housekeeping genes ActB and Gapdh. A list of probes used for this study can be found in table T1.

**Table T1:** List of Taqman assays (mouse and human-specific) used for this study. Probes were purchased from Applied Biosystem.

| *Gene* | *Probe (human)* | *Probe (mouse)* |
| --- | --- | --- |
| Actin | Hs99999903_m1 | Mm_00607939 |
| GAPDH | Hs99999905_m1 | Mm_99999915 |
| Axin2 | Hs00610344_m1 | Mm00443610_m1 |
| P16<sup>Ink4a</sup> | Hs00923894_m1 | Mm_01257348 |
| Bmi1 | Hs00995519_g1 | Mm03053308_g1 |
| Usp16 | Hs01062190_m1 | Mm_00470393 |
| Rspo1 | Hs00543475_m1 | Mm00507077_m1 |
| Rspo2 | Hs00379983_m1 | Mm00555788_m1 |
| Rspo3 | Hs00262176_m1 | Mm00661105_m1 |
| Lgr5 | Hs00969421_m1 | Mm00438890_m1 |

### TTFs, MEFs and human fibroblasts

Mouse primary TTFs were obtained from the tail of wild type, Usp16^+/-^, INK4^−/-^ and INK4^+/-^ as previously described^10^ and used et early passages. Wild-type MEFs were cultured in similar conditions and used at early passages. Human fibroblasts (wild type: CRL-2088, CRL-2076, CTRL-1634 and PCS-201-010; Down’s syndrome: CCL-54, CRL-7090 and CRL-7031) were purchased from ATCC.

#### Wnt activity and stimulation

Mouse and human fibroblasts and basal and luminal mammary cells were treated with recombinant Wnt3a (R&D) at different doses for 16 hours unless otherwise specified. MEFs, Down Syndrome’s and control human fibroblasts were treated in 0. 2% FBS, while mammary cells were sorted, as previously described, and treated with 50ng/ml of Wnt in media supplemented only with Y-27632 and murine recombinant EGF. Wnt activity was measured by Axin2 induction (see above) or luciferase reporter assays. The classical TOPflash (TCF reporter plasmid) and the luciferase βcatenin/TCF reporter 6xKD were kindly provided by Dr. Nusse (Stanford University) and Dr. Boutros (German Cancer Research Center, DKFZ), respectively. As internal control, each reporter was premixed with constitutively expressing Renilla luciferase. Measurements were performed using Centro XS^3^ LB 960 Microplate Luminometer from BERTHOLD Technology.

#### siRNA and lentivirus preparation

USP16 overexpression vector was obtained by subcloning USP16 clone (ATCC) in pCDH-MSCV-GFP vector or pCDH-EF 1A-GFP vector (SBI). TTF and MEF transfection was done by lipofectamine 3000 (Invitrogen). Interfering-RNAs were purchased from Dharmacon. Human fibroblasts were transfected using Lipofectamine RNAi max (Thermos Fisher) following manufacturer’s protocols. Cells were treated with Wnt3a at specified doses 24 hours post-transfection. A list of On-Target PLUS siRNA used for this study can be found in table T2.

**Table T2:**
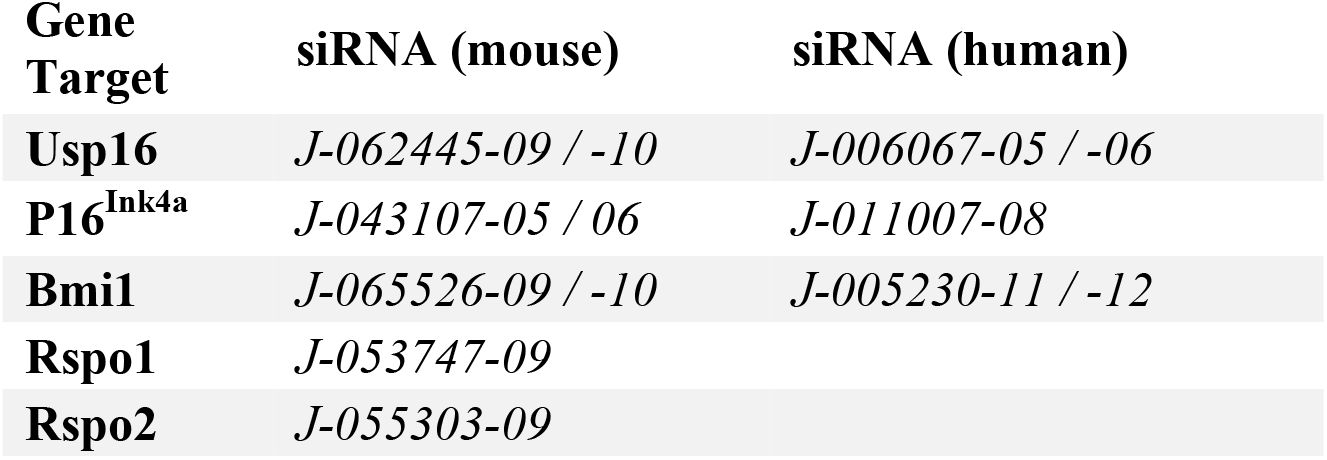
List of siRNA (mouse and human-specific) used for this study. siRNA were purchased from Dharmacon.

#### Co-immunoprecipitation assay

Co-Immunoprecipitation experiments were carried out using the Pierce CO-IP Kit (#26149, Thermo Fisher Scientific) as per manufacturer’s protocol. For β-Catenin and USP16 interaction, MEF P2 were grown in complete medium in a 15cm dish to reach 90% confluence. These were lysed with 1.5 mL ice cold IP Lysis Buffer containing protease and phosphatase inhibitors, for 1 hour at 4°C upon gentle agitation. For the antibody immobilisation step, 10μg of USP16 (Bethyl Laboratories A301-614A) or 10μg of β-Catenin (CST #9562), or as a control 10μg Rabbit IgG, were diluted onto the AminoLink Plus Coupling Resin. The cell lysates were precleared with control agarose resin and co-immunoprecipitation was carried out by adding 1mg of the precleared cell lysate to the antibody immobilised resin, with end over end mixing at 4°C overnight. After elution into 50μL, the sample was analyzed by SDS-PAGE gel and followed by immunoblotting to detect protein-protein interaction.

#### Western blotting

Whole cell lysates were generated by lysis with RIPA buffer, along with protease and phosphatase inhibitor cocktails (Thermofisher Scientific). SDS-PAGE gels were transferred onto PVDF membranes (#IPFL00010, Millipore, Billerica, MA). Membranes were blocked with Li-COR blocking buffer (#927-40000) and subsequently probed with primary antibodies (β-Catenin CST #9562, USP16 Bethyl Labratories A301-614A, Phospho-LRP6 (Ser1490) CST #2568) diluted in 5%BSA/TBST, overnight at 4°C. Incubation with secondary antibodies containing fluorophores at 1:20,000 dilution (IRDye 800CW conjugated goat anti-rabbit #926-32211, LI-COR Biosciences, Lincoln, NE) enabled visualisation on the Odyssey Infrared Imaging System from LI-COR Biosciences.

#### Immunofluorescence microscopy

Cells were seeded on 10mm glass coverslips in full medium and left to adhere for 24 hours to reach 50% confluence. After the appropriate treatments, cells were fixed with ice cold 100% Methanol for 5min at -20°C and then rehydrated thrice in PBS for 5 minutes each. Coverslips were blocked for 30 minutes with 3%BSA/PBS and then incubated with 1:100 dilution of primary antibody (β-Catenin CST #9562) in 1%BSA/PBS for 1 hour in a moist environment. After 3 washes with PBS, cells were incubated with fluorophore labelled secondary antibodies (Alexa Fluorophores Life technologies, Alexa Fluor^®^ 546 A10040) at 1:200 dilution in 1%BSA/PBS for 1 hour. After 3 washes with PBS, the coverslips were mounted using ProLong^®^ Gold Antifade Reagent with DAPI (Thermo Fisher Scientific) as mounting media. Images acquired were quantified for fluorescent intensity on ImageJ and normalized to cell count as measured by DAPI staining.

#### Microarray analyses

Microarray analyses were carried out using the TAC (Transcriptome Analysis Console 4.0) from Applied Biosystem. TAC includes the normalization, probe summarization, and data quality control functions of Expression Console Software. The expression analysis settings were set as fold change <-2 or >2 with a p-value <0.05 using ebayes ANOVA method. For the unsupervised heat map clustering, a gene list including the genes differentially expressed between wt and Ts65Dn (conditional F-test <0.001) was created. A “Wnt signature” gene list, including 413 genes, was derived from the MGI gene ontology annotation (see Suppl. Table 1) (http://www.informatics.jax.org). From this list, a subset of genes differentially expressed between wt and Ts65Dn was chosen for the Wnt-specific clustering (fold change <-1.5 or >1.5 and a conditional F-test <0.05) (see Suppl. Table 2).

## Author contributions

MA and MFC designed the project and wrote the manuscript. MA and BNdR performed most of the experiments. SS and MA performed the breast transplantations assays. VHA helped with managing the mouse colony and with collecting initial data on MMTV-Wnt mammary glands. JA helped with western blots and immunohistochemistry. CH revised the manuscript and oversaw the Down’s syndrome portion of the project.

## Acknowledgments

This study was supported in part by Chan Zuckenberg BioHub, Ludwig Foundation, LumindRDS foundation, DoD postdoctoral fellowship (SS), AIRC and Marie Curie Action (BNdR). BD FACSAriaII was purchased by NIH S10 shared instrumentation grant 1S10RR02933801. Thanks to Samuele Marro, Stanford University, for sharing MEFs.

